# Preserved global cerebral blood flow accounts for youthful processing speed in older adults

**DOI:** 10.1101/665935

**Authors:** Fan Nils Yang, Long Xie, Olga Galli, John A. Detre, David A. Wolk, Hengyi Rao

## Abstract

Preserved cognitive performance is one of the key contributors to successful aging. The processing speed theory and prefrontal executive theory – are competing theories regarding the general causes of cognitive aging. Here, we used a theoretically-driven framework to investigate the neural correlates of older adults with preserved processing speed. Older adults with youth-like processing speed (SuperAgers) were compared with normal aged adults (TypicalAgers) using neuroimaging methods. Global cerebral blood flow (CBF) accounted for approximately 45% of the variance in processing speed, while neither regional CBF nor other structural measures predicted additional variance. In addition, despite having significantly cortical thinning, SuperAgers still shown comparable global CBF levels with young adults. These results support the global mechanism suggested by processing speed theory and indicate that global CBF may serve as a biomarker of cognitive aging.

## 1. Introduction

The world population is aging at an unprecedented rate (Nations, 2015). Individuals are living longer and are expected to survive into their 70s and even 80s on average. Accordingly, it is of utmost socioeconomic importance to promote successful aging and avoid age-related diseases (Eyler et al., 2011; Harada et al., 2013).

Cognitive health is a key contributor to successful aging (Depp and Jeste, 2006; Reichstadt et al., 2007). Cognitive aging refers to a gradual decline in many cognitive abilities (e.g., memory, attention, executive function and processing speed) as humans age (Harada et al., 2013). Despite its importance, the underlying mechanisms of cognitive aging remain obscure.

Various theories and models have been proposed regarding the mechanisms driving cognitive aging. The processing speed theory and the prefrontal executive theory are two competing theories that are among the most influential and empirically tested accounts of age-related cognitive decline (Albinet et al., 2012). The processing speed theory proposed by Salthouse in 1996 suggests that age-related cognitive decline can be accounted for by the *global* mechanism of generalized slowing of cognitive processing (Salthouse, 1996) (see methods for detailed definition of processing speed). The generalized slowing has been associated with reduced global white matter connectivity as indicated by decreased white matter integrity and increase white matter lesion load (Cabeza et al., 2016). Interestingly, other global neural measures not directly related to brain connectivity, such as brain volume and cerebral blood flow (CBF), have also correlated with processing speed (Rabbitt et al., 2007; Rabbitt et al., 2006). On the other hand, the prefrontal executive theory states that *local* structural and functional changes in the frontal cortex lead to a decline in executive function, which in turn produces more general cognitive deficits (West, 1996). Evidence for each these theories has included work demonstrating that after controlling for processing speed or executive function, age-driven differences in high-level cognitive functions are reduced (Anderson et al., 2010; Deary et al., 2010). In addition, behavioral studies have shown that these two theories are not mutually exclusive but share some variance (Albinet et al., 2012). Specifically, executive functions and processing speed each can explain parts of age-related variance on cognition, and they are not mutual exclusive (Albinet et al., 2012).

Examining the neural correlates of older individuals with preserved cognitive functions relative to young adults, i.e. SuperAgers, has been suggested as a promising way to investigate successful cognitive aging and could potentially guide the search for means to improve cognitive decline in older adults (Depp and Jeste, 2006; Eyler et al., 2011; Sun et al., 2016). In this study, we used functional and structural neuroimaging to examine cerebral blood flow, whole brain volume, and cortical thickness in cognitively normal older adults stratified into typical agers (TypicalAgers) and “super” agers (SuperAgers) based on their performance on a simple and well-validated measure of processing speed, i.e. the psychomotor vigilance test (PVT).

It has been well documented that in older adults, big brain-structure size is usually associated with better cognitive performance, especially for frontal regions and executive functions (see Kaup et al. (2011) for a review). Similarly, augmented brain response/activation in frontal regions might severs as a compensatory mechanism in older adults (see Eyler et al. (2011) for a review). However, as Eyler et al. (2011) mentioned, “a simple model of bigger structure → greater brain response → better cognitive performance might not be accurate”. Here we further investigate the potentially different roles of brain structure and function on persevered cognitive function on SuperAgers.

Taken together, the processing speed theory suggested reduced processing speed in TypicalAgers will be associated primarily with global brain measures, such as global CBF or mean cortical thickness of whole brain, while prefrontal executive theory posits that reduction in processing speed will be associated primarily with focal (prefrontal) measures. Alternatively, since these theories are not mutually exclusive, it is also possible that focal and global brain measures each account for unique variance in processing speed. In addition to these theories, previous studies indicated that SuperAgers will likely have bigger brain structure (i.e. thicker gray matter) and greater brain response (i.e. higher CBF during task) than TypicalAgers. More evidences are in need to reveal the underlying mechanism of the “SuperAger phenomenon”.

The current work aim to test these competing hypotheses by investigating both global and focal neural correlates of processing speed in a group of older cognitively normal adults (OG, N = 32). Processing speed was accessed as the mean reaction time (RT) when permorming a well-validated simple reaction time task, i.e. psychomotor vigilance test (PVT) (Basner et al., 2017). Global and regional structural (cortical thickness) and functional (cerebral blood flow (CBF) during performing the PVT task and at rest) neural measurements were extracted from the T1-weighted MRI and arterial spin labeled perfusion MRI (ASL MRI) scans of each subject respectively. In addition, a group of young controls (YC, N = 39) was included in this study serving as a reference group to define cut-off to stratify OG into SuperAgers (N = 15) and TypicalAgers (N = 17) (details described in Section 4.2 and briefly summarized in Figure 1A) as well as to derive data-driven regions of interest (ROI), shown in Figure 1B, to extract local measurements (described in Section 4.4.3). The effect of global and local measurements was compared in terms of discriminating SuperAgers from TypicalAgers as well as predicting processing speed of the older adults (the OG group).

**Figure 1.**
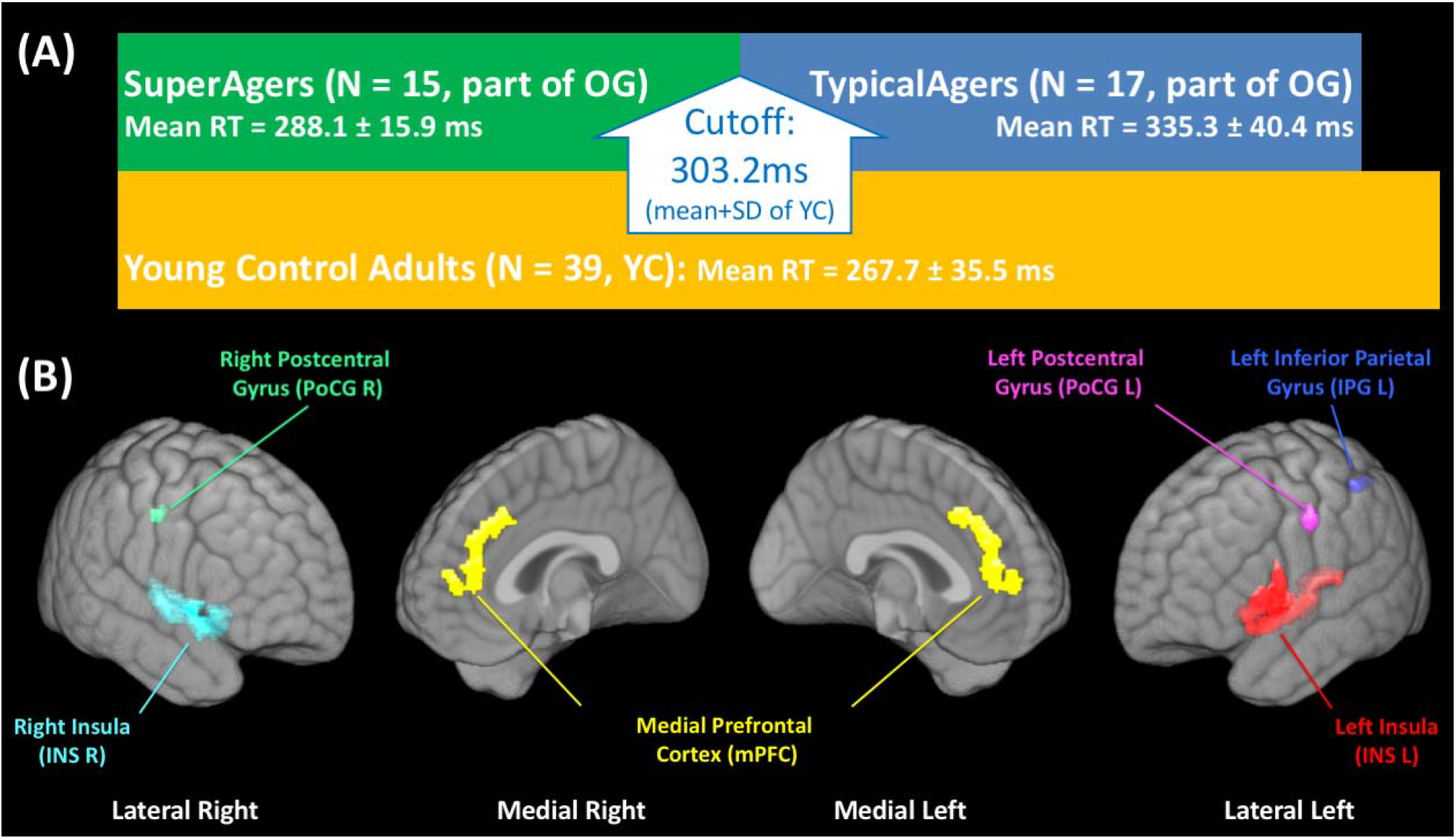
(A) Block diagram showing the assignment of young control (YC), SuperAgers and TypicalAgers. (B) Data driven regions of interest (ROI) in this study. The ROIs were defined in a data-driven manner representing regions with both significant perfusion and structural differences between young controls and older cognitively normal controls (both SuperAgers and TypicalAgers) to locate regions associated with both functional and structural changes due to aging. OG = older cognitive normal adult group, SD = standard deviation.

## 2. Results

### 2.1 Demographic data

The characteristics of YC, TypicalAgers and SuperAgers are shown in Table 1. TypicalAgers and SuperAgers did not differ in sex, age and education. There was no significant difference in Mini-Mental State Examination (MMSE) performance between TypicalAgers and SuperAgers. As expected, there was no significant difference in mean reaction time (RT) between SuperAgers and YC.

**Table 1.**
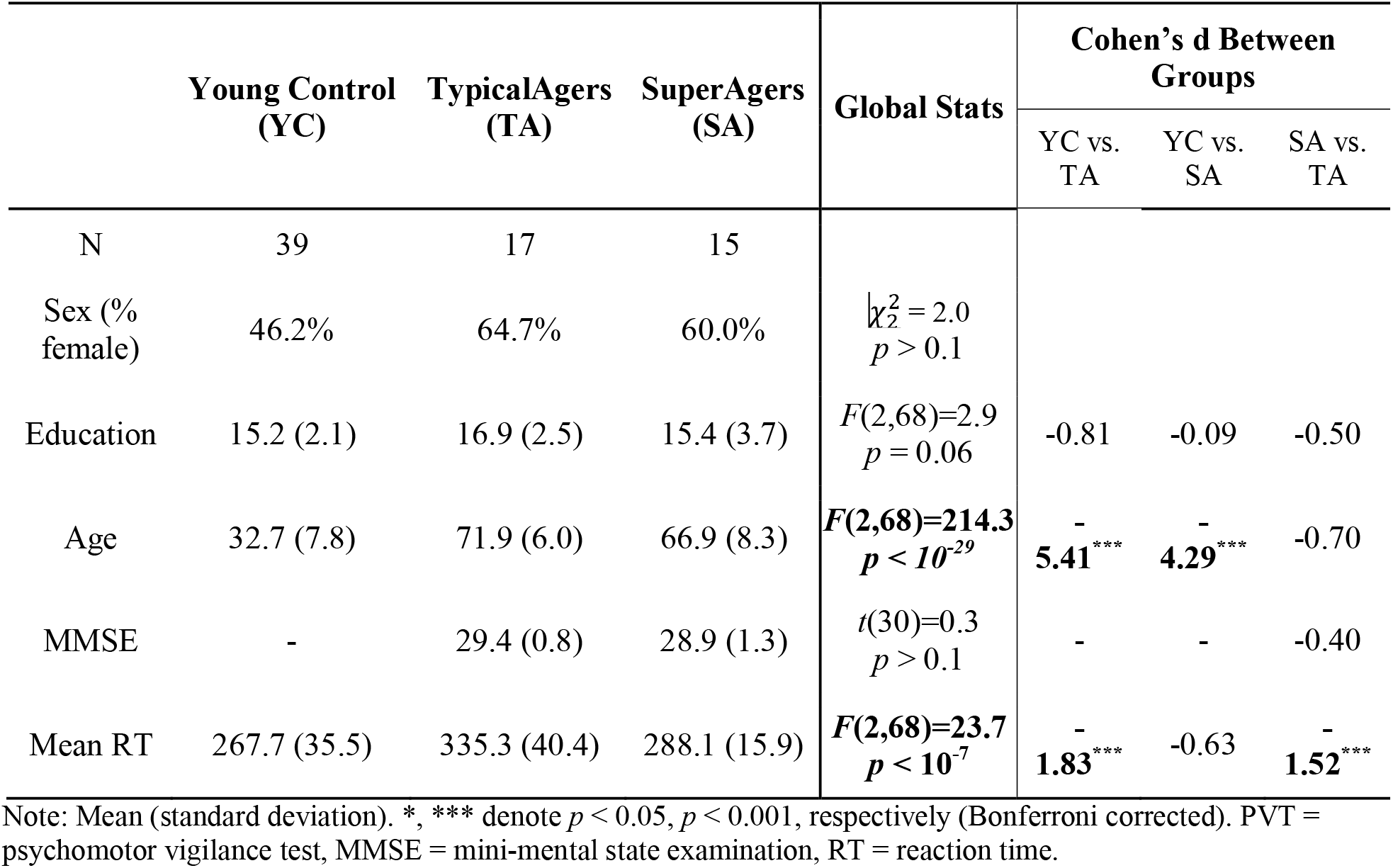
Demographics, MMSE and mean PVT reaction time of young controls, TypicalAgers and SuperAgers.

### 2.2 Discriminating SuperAgers and TypicalAgers

To investigate potential global effects, the whole-brain mean resting CBF, whole-brain task CBF, whole brain mean cortical thickness, cortical gray matter and white matter volume of each subject were extracted. Analysis of covariance (ANCOVA) covaried for age, sex and education revealed significant group differences in functional and structural global measurements in the three groups (Table 2), including differences in whole-brain CBF during PVT task (*F*(2,68) = 21.4, *p* <10^-7^) and at rest (*F*(2,68) = 18.5, *p* <10^-6^), mean cortical thickness (*F*(2,68) = 100.9, *p* <10^-20^), as well as cortical gray matter volume (*F*(2,68) = 85.8, *p* <10^-18^). Post-hoc analyses results, reported in Table 2, demonstrated that the major differences between SuperAgers and TypicalAgers were found in whole-brain task and rest CBF (all *p* < 0.001 corrected with large Cohen’s d). Also, SuperAgers had significantly more cortical gray matter than TypicalAgers (*p* < 0.05 corrected with relatively small Cohen’s d). It is also worth noting that no differences in whole-brain task or rest CBF were found between YC and SuperAgers (all *p* > 0.05), while this was not the case between YC and TypicalAgers (all *p* < 0.001 corrected).

**Table 2.** Global and structural measurements of Young Controls, TypicalAgers and SuperAgers.

|  | Young Control (YC) | Typical Agers (TA) | Super Agers (SA) | ANOVA Global Stats | Cohen's d Between Groups |  |  |
| --- | --- | --- | --- | --- | --- | --- | --- |
|  |  |  |  |  | YC vs. TA | YC vs. SA | SA vs. TA |
| N | 39 | 17 | 15 |  |  |  |  |
| Whole-Brain Task CBF (ml/100g tissue/min) | 63.8 (12.9) | 42.2 (9.5) | 56.3 (8.4) | $F(2,68)=21.4$<br>$p < 10^{-7}$ | **1.80\*\*\*** | 0.62 | **1.58\*\*\*** |
| Whole-Brain Resting CBF (ml/100g tissue/min) | 62.9 (13.6) | 42.1 (9.7) | 56.6 (8.0) | $F(2,68)=18.5$<br>$p < 10^{-6}$ | **1.66\*\*\*** | 0.51 | **1.62\*\*\*** |
| Cortical Gray Matter Volume (1000 mm<sup>3</sup>) | 551.4 (21.6) | 467.4 (29.1) | 492.0 (22.6) | $F(2,68)=85.8$<br>$p < 10^{-18}$ | **3.49\*\*\*** | **2.71\*\*\*** | **0.94\*** |
| White Matter Volume (1000 mm<sup>3</sup>) | 451.1 (15.2) | 442.0 (24.4) | 447.2 (14.2) | $F(2,68)=1.6$<br>$p > 0.1$ | 0.27 | 0.44 | 0.26 |
| Mean Cortical Thickness (mm) | 2.18 (0.15) | 1.69 (0.07) | 1.79 (0.14) | $F(2,68)=100.9$<br>$p < 10^{-20}$ | **3.81\*\*\*** | **2.66\*\*\*** | 0.93 |

Note: Mean (standard deviation). \*, \*\*\* denote $p < 0.05, p < 0.001$ , respectively (Bonferroni corrected). CBF = cerebral blood flow.

In addition to global measurements, the mean resting CBF, task CBF and cortical thickness of each ROI shown in Figure 1B were extracted as local measurements. Significant group differences, tested by ANCOVA accounting for age, sex, and education, between SuperAgers and TypicalAgers were seen in CBF measurements during the PVT task and at rest (see Figure 2). Midline structures in the frontal lobe, (i.e. mPFC) exhibited group differences at a trend level in cortical thickness (*t*(30) = 3.2, *p* = 0.003, *p* >0.05 after correction).

**Figure 2.**
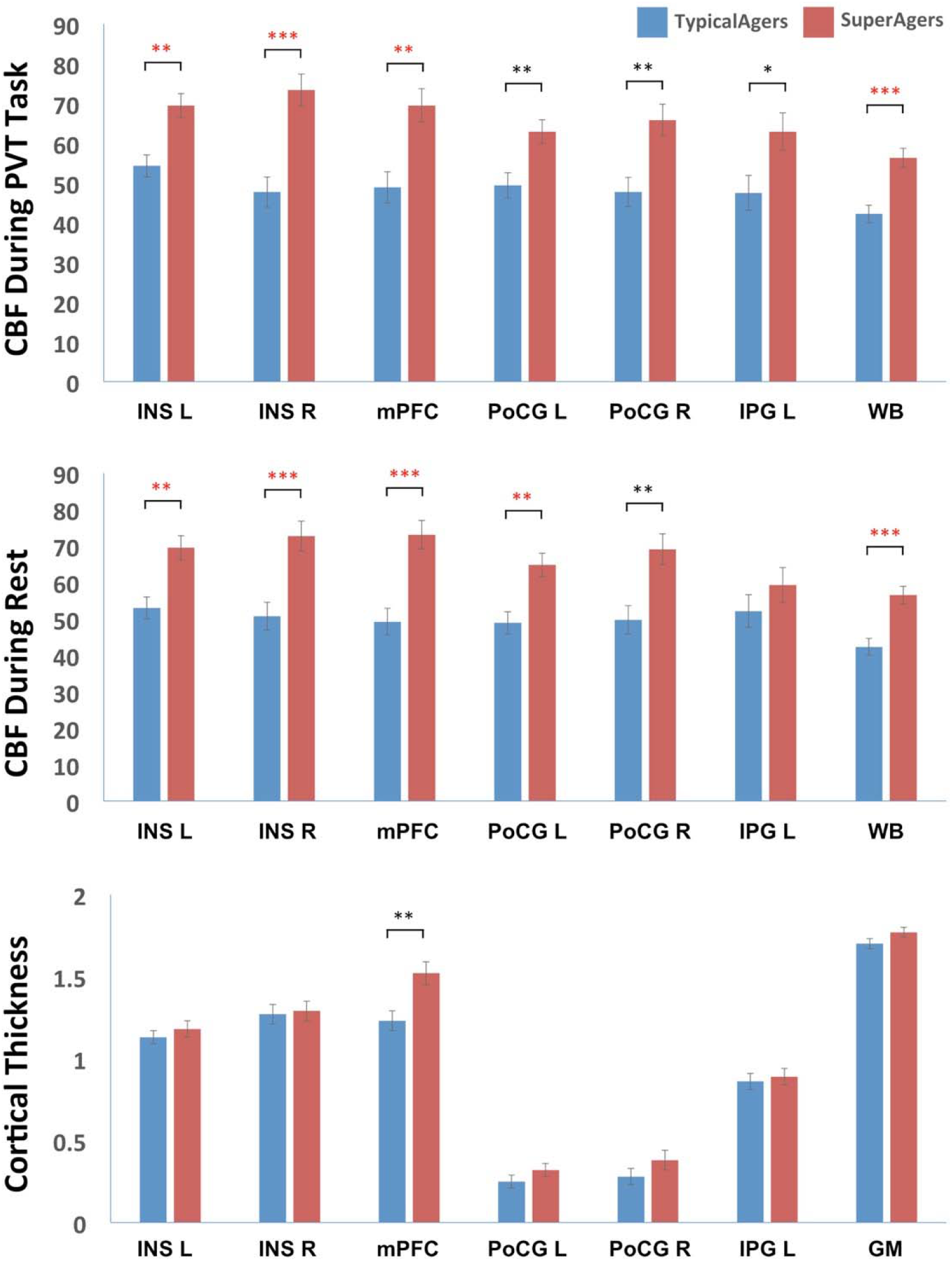
Regional and global cerebral blood flow (CBF) and cortical thickness measurements, adjusted for age, sex and education. Abbreviations: INS = insula, mPFC = medial prefrontal cortex, PoCG = postcentral gyrus, IPG = inferior parietal gyrus, WB = whole brain, GM = gray matter, L = left, R = right. *, **, *** denote *p* < 0.05, *p* < 0.01, *p* < 0.001, respectively. Red stars indicate *p* values survived Bonferroni corrections (*p* < 0.05/21).

To further illustrate the differences between the three groups, bar plots of each of the regional or global functional and structural measurements for each of the three groups without adjustment for covariates are shown in Supplementary Figure S3 in Supplement C. Qualitatively, we observed that regional resting and task CBF in SuperAgers were more similar to YC compared to that of the TypicalAgers. However, the amount of cortical thinning of both SuperAgers and TypicalAgers is similarly significantly lower than YC. This may indicate that, compared to Typical Agers, brain function of SuperAgers (as reflected by CBF measures) is more preserved even in the context of similar cortical atrophy relatively young adults.

### 2.3 Correlations between CBF and mean reaction time

CBF during the PVT task and at rest were significantly correlated with mean RT (Table 3). However, none of these correlations remained significant when including the corresponding global measurement (i.e global measure of task CBF, rest CBF and thickness for ROI-based task CBF, rest CBF and thickness measures respectively) as a covariate. Figure 3 shows a scatter plot of age, sex and education-adjusted mean RT and global CBF during the PVT task, which was the most predictive measurement for processing speed (*r* = 0.71, *p* < 0.001). Cortical thickness measures in ROIs and total cortical gray matter were not predictive of mean RT. The two-step hierarchical linear regression further demonstrated that only whole-brain task CBF (*β* = −2.5, *R^2^* change = 0.451, *p* < 0.001) was included in the most predictive model (N = 32, *F* = 7.5, *R^2^* = 0.526, *p* < 0.001). Structural measurements provided no additional information.

**Table 3.**
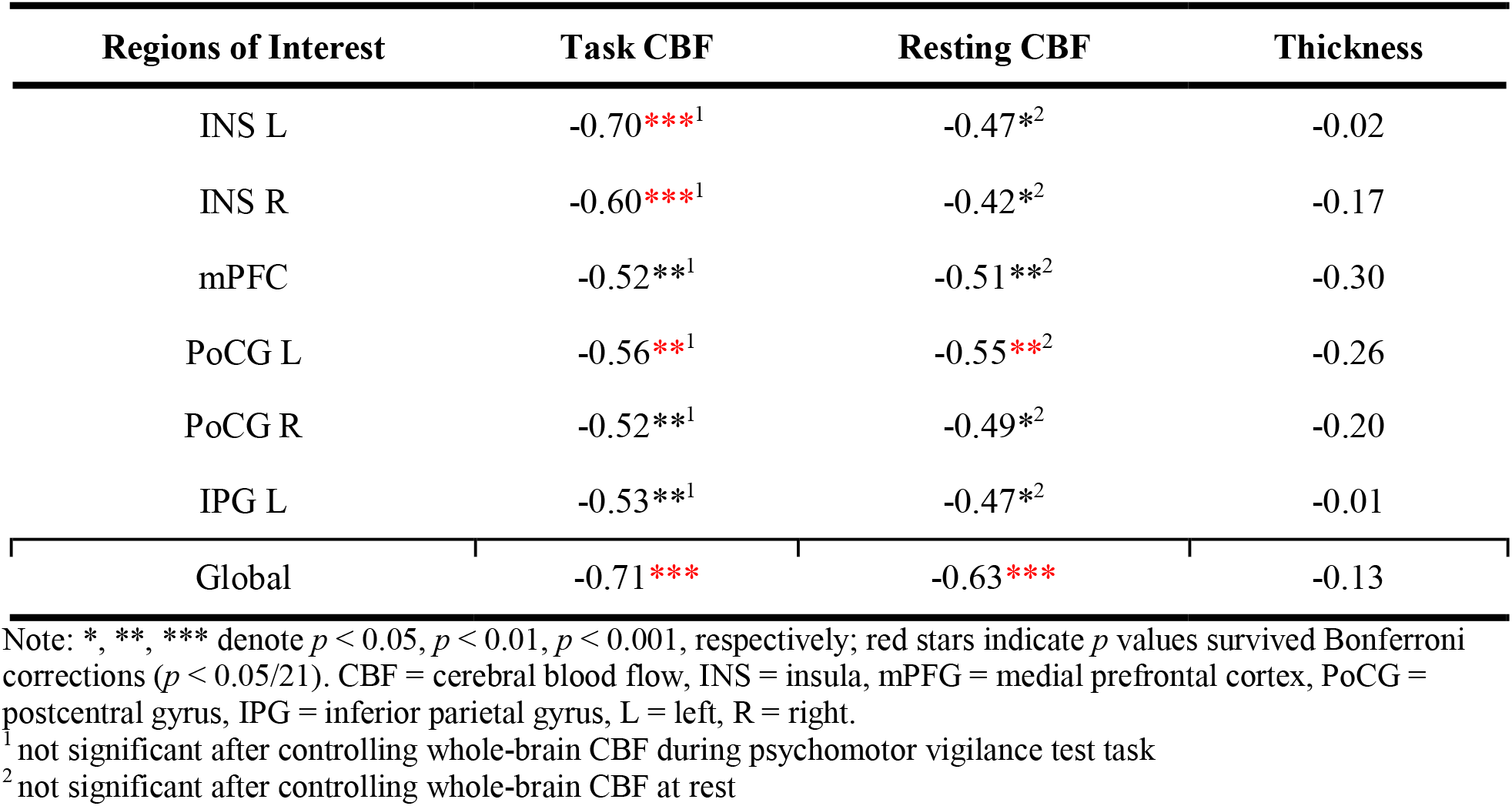
Partial Pearson correlations between global and ROI-based functional and structural measurements and mean PVT reaction time, with age, sex and education as covariates.

**Figure 3.**
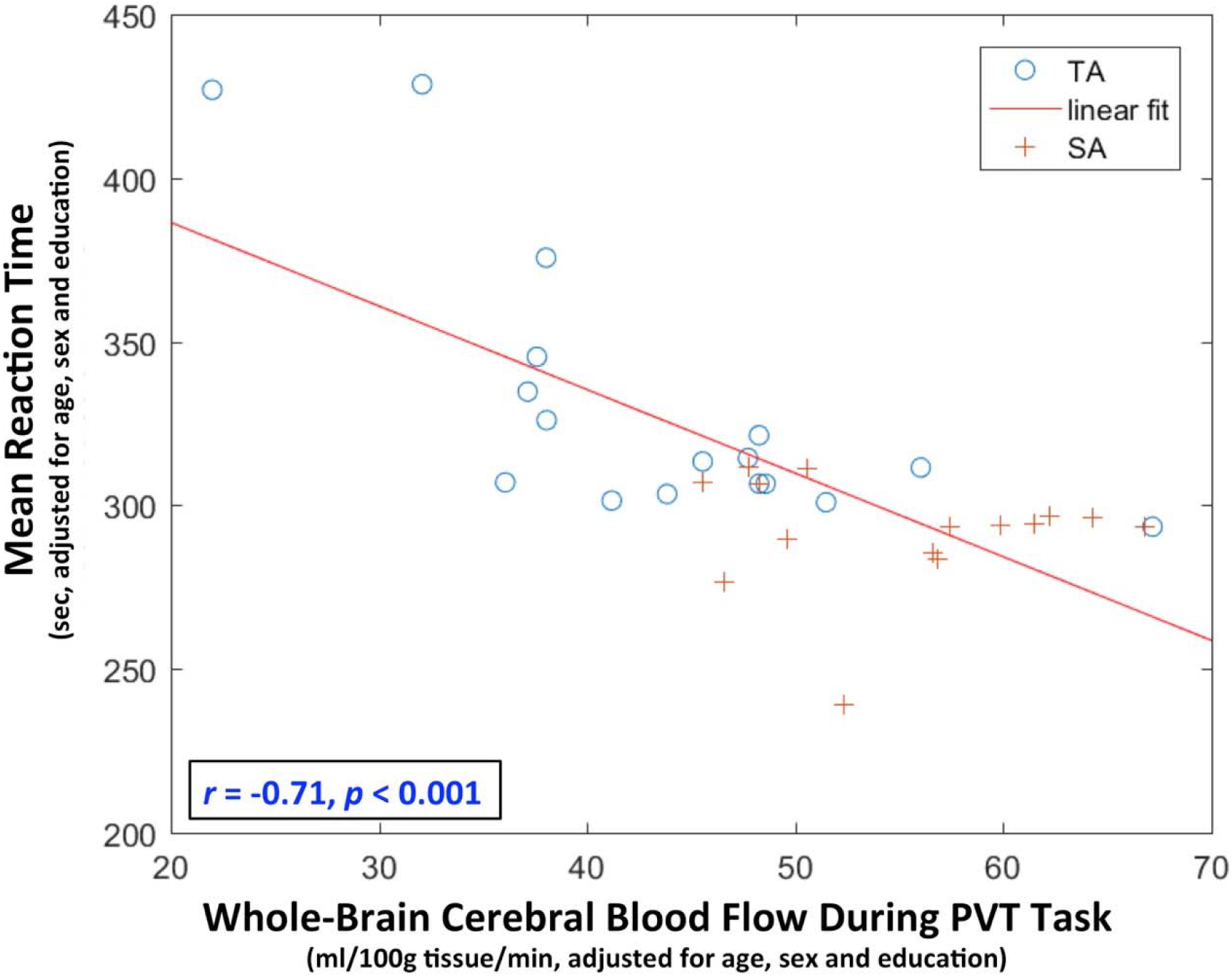
Scatter plot of the whole-brain cerebral blood flow during the psychomotor vigilance test (PVT) and mean reaction time, adjusted for age, sex and education, of TypicalAgers (TA) and SuperAgers (SA).

Voxel-wise partial correlation analyses between resting and task CBF and mean RT (Supplement D) demonstrated that both resting and task CBF across a majority of brain regions is significantly correlated with mean RT (Supplementary Figure S4).

## 3. Discussion

The principal purpose here was to examine the neural correlates of persevered processing speed in SuperAgers. The present findings demonstrate that preserved processing speed in SuperAgers is associated with global CBF both at rest and more so during PVT task performance, rather than with regional CBF. SuperAgers showed global rest and task CBF values that were not significantly different from young controls, despite having significantly thinner cortical thickness and lower regional CBF in frontal regions. This effect was observed even after controlling for demographic factors such as age, gender, and education, as well as other brain measurements such as whole brain volume and cortical thickness.

### 3.1 The definition of SuperAgers

Previously, the term SuperAgers has been used to refer to older adults with youthful memory abilities at normative performance for young adults on delayed free recall tests (Gefen et al., 2014; Harrison et al., 2017; Sun et al., 2016). We extended the definition to older adults that have a comparable performance with young adults on a simple reaction time task because some studies have shown that when controlling for processing speed, age differences in memory may be largely reduced or even eliminated (Bryan and Luszcz, 1996; Lee et al., 2012). Furthermore, processing speed is thought to be one of the behavioral measures most sensitive to age (Cabeza et al., 2016).

### 3.2 Global CBF, but not regional CBF, correlated with processing speed in older adults

Although previous studies have shown that total resting CBF is positively correlated with processing speed in older adults (Poels et al., 2008; Rabbitt et al., 2007; Rabbitt et al., 2006), this relationship has not been well-understood. For example, by using phase-contrast MRI, these studies did not assess regional CBF. It is possible that the use of total CBF masked regional relationships, as Steffener et al. (2013) showed that distributed regional CBF correlated with processing speed measures in older adults. The current study extended previous studies by demonstrating that the correlation between regional CBF and processing speed in older adults might be fully driven by the global CBF. In addition, the global CBF during task explained more variance in processing speed as compared to the global resting CBF, suggesting that task CBF might be more sensitive to the processing speed in older adults and should be considered as one biomarker for processing speed in the future. This notion is in line with previous studies, which demonstrated that task-state CBF may be more associated with cognitive decline (Xie et al., 2018; Xie et al., 2016).

The fact that global, but not regional CBF accounted for the processing speed in older adults provides an additional neural evident for the processing speed theory. According to this theory, degraded cognitive functions in old age is due to the global mechanism of the slowed processing speed, which limits the ability to simultaneously process a certain amount of information needed for higher level cognitive functions (Salthouse, 1996). This generalized slowing is associated with the decreased efficiency of interregional communication, which might be caused by decreased white matter integrity (Cabeza et al., 2016). Indeed, decreased white matter integrity of the whole brain is associated with this generalized slowing (Cabeza et al., 2016; Gunning - Dixon et al., 2009; Penke et al., 2010). The current study extended this notion to the CBF. Recent studies have shown that there is a significant relationship between CBF and white matter integrity (Brickman et al., 2009; Chen et al., 2013). It is possible that global CBF and white matter integrity of whole brain both plays an important role on supporting the efficiency of interregional communication throughout the brain. Future studies should determine whether CBF and white matter integrity account for unique portions of variance in processing speed. Another possible explanation regarding this CBF-speed association is that as CBF is tightly coupled with brain metabolism (Raichle, 1998), reduced global CBF and brain metabolism likely reflect global neuronal dysfunction. Therefore, reduced CBF is leading to less efficient brain and decreased processing speed. This hypothesis needs to be tested in future studies.

### 3.3 Neither functional or structural changes in specific regions (e.g. frontal lobe) accounted for processing speed in older adults

The rationale for prefrontal executive theory is as follow, 1) executive functions are a set of high-order functions that are necessary for the cognitive control and coordination of fundamental cognitive operations, 2) the frontal lobe is a key region for executive functions, 3) both executive functions and the frontal lobe are extremely sensitive to the effects of normal aging (Phillips and Henry, 2008). Consistent with this rationale, both large age-related CBF reductions and cortical thinning was were found in prefrontal regions and bilateral insula in both groups of older adults. In addition, large CBF and cortical thickness difference in medial prefrontal cortices were found between SuperAgers and TypicalAgers, suggesting the executive function might be different between these two groups as well. Nonetheless, neither thickness within medial prefrontal cortex nor other cortical thickness measurement were correlated with processing speed.

In contrast to previous studies showing that CBF and brain volume each account for unique variance in processing speed (Rabbitt et al., 2007; Rabbitt et al., 2006; Steffener et al., 2013), or that brain volume mediates the association between CBF and processing speed (Poels et al., 2008), the present study found no significant differences between SuperAgers and TypicalAgers in white matter volume, and relatively small difference in gray matter volume. One possible explanation is that age itself has a large effect on CBF, brain volume, processing speed, and other brain measures (see Cabeza et al. (2016) for a review). This possible confound was minimized by focusing on comparisons between SuperAgers and age-matched TypicalAgers. Step-wise regression results further confirmed that global task related CBF was the only significant predictor of PVT performance in older individuals, accounting for 45.1% of the variance in mean response time.

Taken together, although regional structural and functional differences were found between SuperAgers and TypicalAgers, it is the global functional measure CBF that explained the most variance in processing speed. In addition, as indicated in Supplementary Figure S3, it is surprising that with similar amount of significant cortical thinning as TypicalAgers (much lower compared to young adults), SuperAgers are able to preserve youth-like processing speed and to maintain youthful global CBF. These results provide further evidence that a simplified model of “bigger brain structure, better cognitive performance” may not be sufficient in explaining successful cognitive aging (Eyler et al., 2011). Instead, the robust relationship between global CBF and processing speed suggested that global CBF might be considered as a potential biomarker for successful cognitive aging.

### 3.4 Limitations

There are several limitations to this study. First, as we did not directly test executive function in the current study, it is possible that executive functions were not different between SuperAgers and TypicalAgers, and hence here the focal regions did not contribute to the processing speed difference between the two groups. However, previous studies have shown that the majority of the age-related variance in executive functions is shared with processing speed (Albinet et al., 2012; Luszcz, 2011). In addition, substantial differences were found in prefrontal regions, in terms of both cortical thickness and CBF measures, between these two groups, suggesting executive functions differences might be also differ between SuperAgers and TypicalAgers. Second, it remains an open question whether the SuperAgers in the present sample were also top performers in their youth. Based on our results, it is possible that young adults with higher processing speed performance were resilient against age-related CBF decline and remained top performers as they aged. Longitudinal studies are still needed to test this possibility. Third, the current study used only one processing speed measure, which raises a question of whether these results are task-specific. Salthouse (2000) found that all six speed measurements using different tasks, including reaction time, are highly intercorrelated and that age-related effects on an individual speed measure can be largely explained by other speed measures. Thus, it is likely that our results can be repeated by using other measures of processing speed.

### 3.5 Future directions

Future studies may utilize larger datasets to investigate the relationship between global CBF and age-related decline in a broader range of cognitive domains. Interestingly, a recent study found that global CBF can predict older adults’ general cognitive function 4 years later (De Vis et al., 2018). This finding further supports our conclusion that global CBF may serve as a biomarker of processing speed, and therefore as a biomarker of cognitive aging (as suggested by the processing speed theory).

Previous studies have shown that slowed reaction times are associated with an elevated risk of future cognitive disorders (Cherbuin et al., 2010; Kochan et al., 2016). Given the robust relationship between global CBF and reaction time, future studies may wish to examine whether reduced global CBF is one of several key predictors of transition from healthy aging to dementia. Further investigation of the neural mechanisms underlying the SuperAger phenomenon may allow us to design effective interventions to promote successful aging.

## 4. Methods and materials

### 4.1 Study design

The current study is an observational study. Data from eighty-three healthy adults aggregated across two different study cohorts are included in the analyses. Forty-three young adults (21-50 years old) comprised the young control group (YC). These subjects were recruited in response to study advertisements, as part of a sleep deprivation study (Fang et al., 2015). The remaining 40 subjects (57-85 years old), referred to as the older cognitively normal group (OG), were recruited from the Penn Memory Center as part of a study of prodromal Alzheimer’s disease (Xie et al., 2016). All subjects had no history of clinical stroke, significant traumatic brain injury, alcohol or drug abuse/dependence, or any other medical or psychiatric condition thought to significantly impact cognition. The study was approved by the Institutional Review Board of the University of Pennsylvania.

### 4.2 Processing speed task: psychomotor vigilance Test

By definition, processing speed refers to the speed of motor responses and the speed with which cognitive operations can be executed (Harada et al., 2013; Salthouse, 1996). Processing speed tends to be affected earlier in lifespan, as compared to other cognitive abilities, like episodic memory, reasoning, and spatial ability (Hedden and Gabrieli, 2004; Salthouse, 1996; Schaie, 1996).

Here, a well-validated simple reaction time task, PVT, was used. PVT has been shown to be highly reliable, free of learning/practice effect, and uncontaminated by aptitude (Basner et al., 2017), making it suitable for assessing processing speed in older populations. During the PVT, participants are instructed to maintain their attention on a red-outlined rectangular area located in the center of a dark screen and to respond (button press) as fast as they can whenever a yellow millisecond counter appears inside the rectangle. The millisecond counter stops after participants’ action and remains for another one second to allow participants to see their reaction time (RT). Button presses when the millisecond counter did not appear are counted as false alarms, whereas failure to respond within 30 seconds leads to a time out. Participants are instructed to respond as quickly as possible while maintaining accuracy. The PVT was administered during arterial spin labeled perfusion MRI (ASL MRI) scanning. The reaction time measures, excluding those from time out trials, were averaged to compute mean RT for each subject, which serves as a measure of the subject’s processing speed.

The OG group was divided into TypicalAgers and SuperAgers based on their mean RT during the PVT. The cutoff was determined as one standard deviation above the mean (slower responses) of the YC group, which was 303.2 milliseconds. OG individuals with a mean RT within one standard deviation of the young control group were categorized as SuperAgers, while those with a mean RT greater than 303.2ms were categorized as TypicalAgers. The group assignment was summarized in Figure 1A.

### 4.3 MRI acquisition

All imaging was performed on a 3T Siemens Trio MRI scanner (Erlangen, Germany) equipped with either a product eight-channel or thirty-two-channel array coil. The scans of YC and OG were acquired following two similar protocols. In both protocols, high-resolution structural images were acquired with 3D-MPRAGE (Mugler III and Brookeman, 1990) at 1 mm^3^ isotropic resolution (TI = 950 ms, TE = 3 ms, TR = 1620 ms). In addition, a pseudocontinuous ASL (pCASL) (Dai et al., 2008) with a 2D gradient-echo echo planar imaging (GR-EPI) readout was used in both protocols to measure regional cerebral blood flow (CBF). All participants were scanned during a ‘resting’ and a ‘task’ sequence.

Acquisition parameters for the pCASL sequence in the protocol for OG were: TR/TE/FA = 4 s/19 ms/ 90°, 6 mm slice thickness, 1 mm inter-slice gap, 18 slices acquired in ascending order, 3.5 × 3.5 × 7 mm^3^ resolution. Arterial spin labeling was implemented with mean Gz of 0.6 mT/m and 1640 Hanning window shaped RF pulses for a total labeling duration of 1.5 seconds. The labeling plane was positioned 80 or 90 mm below the center of the imaging region and post-labeling delay was set to 1.5 seconds. The ‘resting’ and ‘task’ sequences lasted ~6 min with 45 pairs of label-control scans for signal averaging. Due to technical issues or subject fatigue, only 37 pairs of ‘task’ scan for one subject were acquired, but data quality was sufficient for inclusion in the analyses.

The pCASL sequence in the protocol for YC differed from that of the OG in the following aspects: (1) The post-labeling delay of the arterial spin labeling was set to 1.0 second to account for reduced transit time in YC; (2) Due to a change of protocol during the study, the ASL data was acquired in two slice thickness: either 4.8 mm (30 slices) or 6 mm (20 slices); (3) 30 pairs of label-control scans were acquired for the ‘resting’ sequence; (4) the ‘task’ sequence acquired 75 pairs of scans. Due to potential subject fatigue during ‘task’ ASL MRI (Lim et al., 2010), different task durations may introduce bias. To avoid this, the ‘task’ sequence of the YC was truncated and only the first 45 pairs of scans were analyzed. This was not necessary for the ‘resting’ CBF measurement.

### 4.3 Neuroimaging data processing

#### 4.3.1 Tissue segmentation and CBF quantification

Statistical Parametric Mapping 8 (SPM 8, Wellcome Department of Cognitive Neurology, UK) and ASLtbx (a SPM add-on toolbox) were used to perform tissue segmentation and to quantify regional cerebral blood flow (CBF) from the ASL MRI scans, including the following steps: (1) realignment and averaging of ASL time series to correct for head motion and to generate a mean EPI image, (2) rigid registration of the mean EPI image to the anatomical image, (3) transformation of each frame of the ASL time series using the image generated in the previous step followed by smoothing in space with a 3-dimensional 4 mm full width at half maximum (FWHM) Gaussian kernel, (4) tissue segmentation of the structural image using the pipeline in SPM8 to generate gray matter (GM), white matter (WM) and cerebrospinal fluid (CSF) probability maps, which are then smoothed using a 3-dimensional FWHM Gaussian kernel and resampled to the space of the registered mean EPI image, (5) generation of perfusion-weighted time series using pairwise subtraction of the label and control images, (6) application of the modified single compartment continuous ASL perfusion model (Wang et al., 2003) to the perfusion-weighted time series to derive an absolute CBF image series, (7) application of the Structural Correlation-based Outlier Rejection (SCORE) (Dolui et al., 2017) to the CBF image series to perform denoising and generate a “cleaned” CBF image, (8) adjustment of the CBF signal at each voxel of the “cleaned” CBF image by dividing with the GM probability plus 0.4 times WM probability at the corresponding voxel to correct for partial volume effect (results did not change without partial volume effect, see Supplement A), (9) normalization to the 2×2×2 mm^3^ Montreal Neurological Institute (MNI) template using the DARTEL algorithm (Ashburner, 2007).

#### 4.3.2 Cortical thickness map estimation

A diffeomorphic registration based cortical thickness analysis pipeline (Das et al., 2009) available in Advanced Normalization Tools (ANTs) was applied to the structural MRI scan of each subject to derive a voxel-wise cortical thickness map. The thickness maps were smoothed using a 4 mm FWHM Gaussian kernel and normalized to the MNI space (1×1×1 mm^3^).

#### 4.3.3 Quality control

Quality control was performed by visual inspection. Subjects with CBF maps with extensive non-physiological negative or positive CBF clusters (likely caused by instability of spin labeling, subject motion, or MRI artifacts) were identified and excluded from the study. The SPM DARTEL pipeline failed on one individual in the older adult group. In total, data for four YC and four OG subjects were excluded. In addition, the four oldest TypicalAgers were excluded to generate an age-matched sample with the SuperAgers, yielding 39 YC, 15 SuperAgers and 17 TypicalAgers (i.e. 32 OG) for the final analysis.

### 4.4 Statistical analysis

Statistical analyses were performed using standard methods in SPSS 23.0 (Chicago, IL), MATLAB 2014a (Math Works, Natick, MA) and FSL 5.0.5. All statistical analyses are two-tailed.

#### 4.4.1 Analysis of demographic data

Contingency χ^2^ testing, and analysis of variance (ANOVA) with post-hoc comparisons were used to test group differences among YC, TypicalAgers and SuperAgers. Bonferroni correction was applied to adjust for multiple comparisons.

#### 4.4.2 Analysis of global measurements

To investigate potential global effects, the whole-brain mean resting CBF, whole-brain task CBF, whole brain mean cortical thickness, cortical gray matter and white matter volume of each subject were extracted. ANOVA analysis with post-hoc comparisons and Bonferroni correction was used to investigate group differences and perform pairwise group comparisons between YC, TypicalAgers and SuperAgers.

#### 4.4.3 Analysis of local measurements

A region of interest (ROI) analysis was also used to analyze neuroimaging measurements. The ROIs were defined in a data-driven manner representing regions with both significant perfusion and structural differences between YC and OG to locate regions associated with both functional and structural changes due to aging. Voxel-wise independent two sample t-tests were performed using the normalized resting CBF maps and thickness maps separately using the “Randomize” package (Winkler et al., 2014), with family-wise error rate (FWE) correction for multiple comparisons. As cortical thickness was only defined in cortex, the analysis for the thickness maps was limited to voxels with cortical thickness greater than 0.1mm in all the subjects. Regions with corrected *p* <= 0.001 in both analyses were defined as ROIs. Individual statistical maps of resting CBF and cortical thickness together with the ROIs are shown in Supplement B. ROIs with cluster sizes smaller than 320 mm^3^ (corresponding to 40 voxels in the 2×2×2 mm^3^ MNI template) were excluded. In total, six ROIs were generated (Figure 1B), and included the bilateral insula (INS L/R), bilateral postcentral gyrus (PoCG L/R), medial portion of the prefrontal gyrus (mPFC), and left inferior parietal gyrus (IPG L). In each ROI, mean resting CBF, task CBF, and cortical thickness were extracted for each subject.

For each global or ROI-based measurement, an analysis of covariance (ANCOVA), with age, sex, and education as covariates was performed to examine the difference between TypicalAgers and SuperAgers. Within TypicalAgers and SuperAgers, ROI-based partial linear correlation analyses controlling for age, sex and education, was performed between mean RT and neuronal measurements. Bonferroni correction was applied to correct for multiple comparisons. To investigate whether measurements of the regional ROIs provide additional predictive value beyond the global measurements, the partial correlation analysis was repeated with the corresponding global measurement as an additional covariate.

#### 4.4.4 Hierarchical linear regression analysis for functional and structural measurements

To further investigate whether structural measurements provide complementary information in predicting mean RT, a two-step, hierarchical linear regression was performed with age, sex, and education entered in the first step, and with whole-brain task and resting CBF and structural measurements, including ROI-based and whole-brain mean cortical thickness, gray matter volume, and white matter volume in the second step.

## Author contributions

Study concept and design: R.H., D.A.W., J.A.D.

Data analysis: F.N.Y., L.X.

Drafting manuscript: F.N.Y., L.X., O.G.

Approval of the final version of the manuscript: All authors.

## Supplementary Materials

### A. Statistical analysis results using CBF measurements without corrected for partial volume effect

All of the statistical analyses were repeated using CBF measurements without correction for a partial volume effect. The results, shown in Table S1, Table S2, and Figure S1 are similar to those with partial volume correction. Similarly, the two-step, hierarchical linear regression using the uncorrected CBF measurements generated similar results with only whole-brain task CBF (*β* = −3.1, *R^2^* change = 0.457, *p* < 0.001) selected in the most predictive model (N = 32, *F* = 7.7, *R^2^* = 0.532, *p* < 0.001).

**Table S1.**
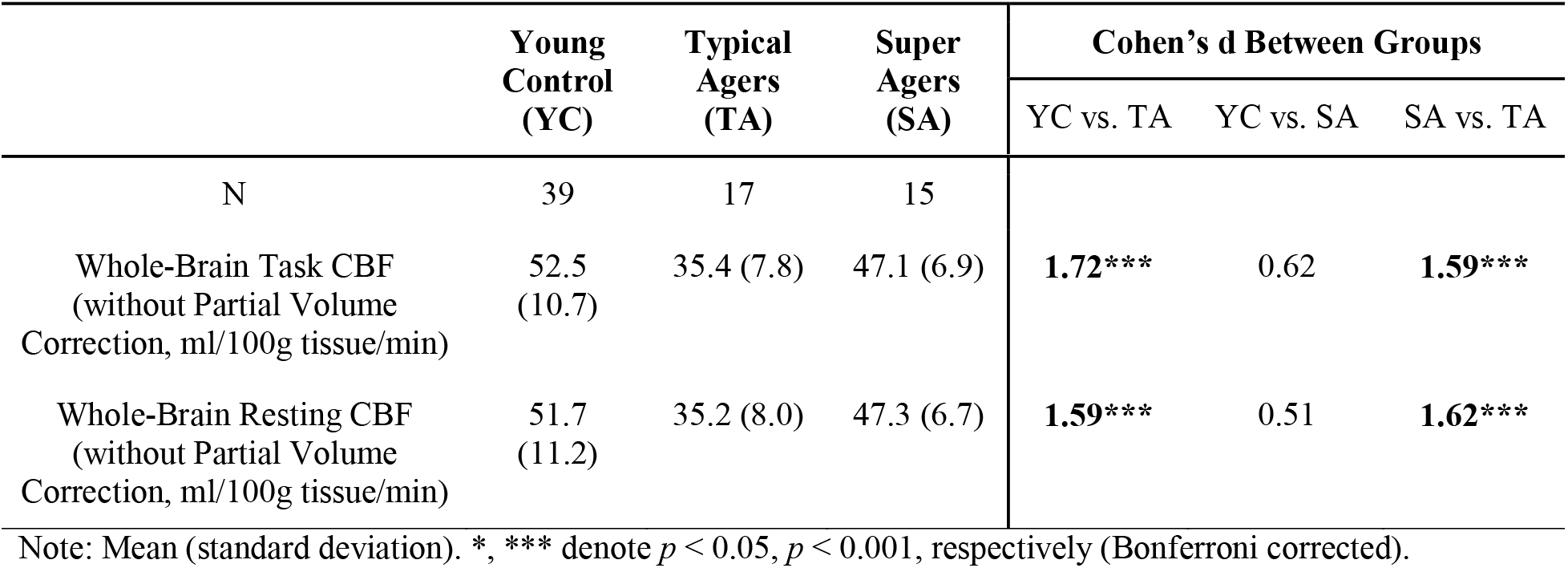
Global functional measurements in Young Control, TypicalAgers and SuperAgers without correction for partial volume effect.

**Table S2.**
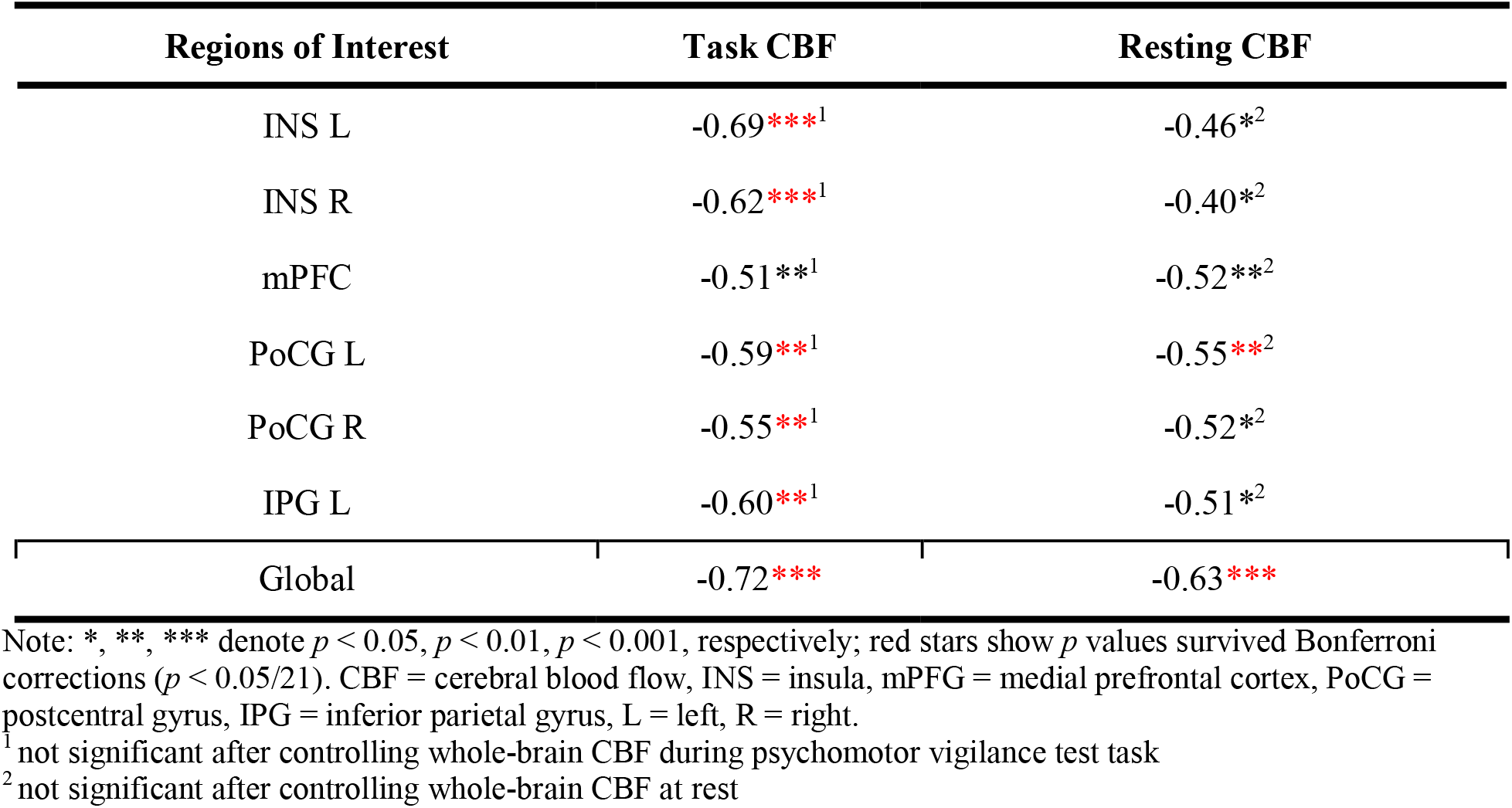
Partial Pearson correlations between global and ROI-based functional measurements without correction for partial volume effect and mean reaction time, with age, sex and education as covariates.

**Figure S1.**
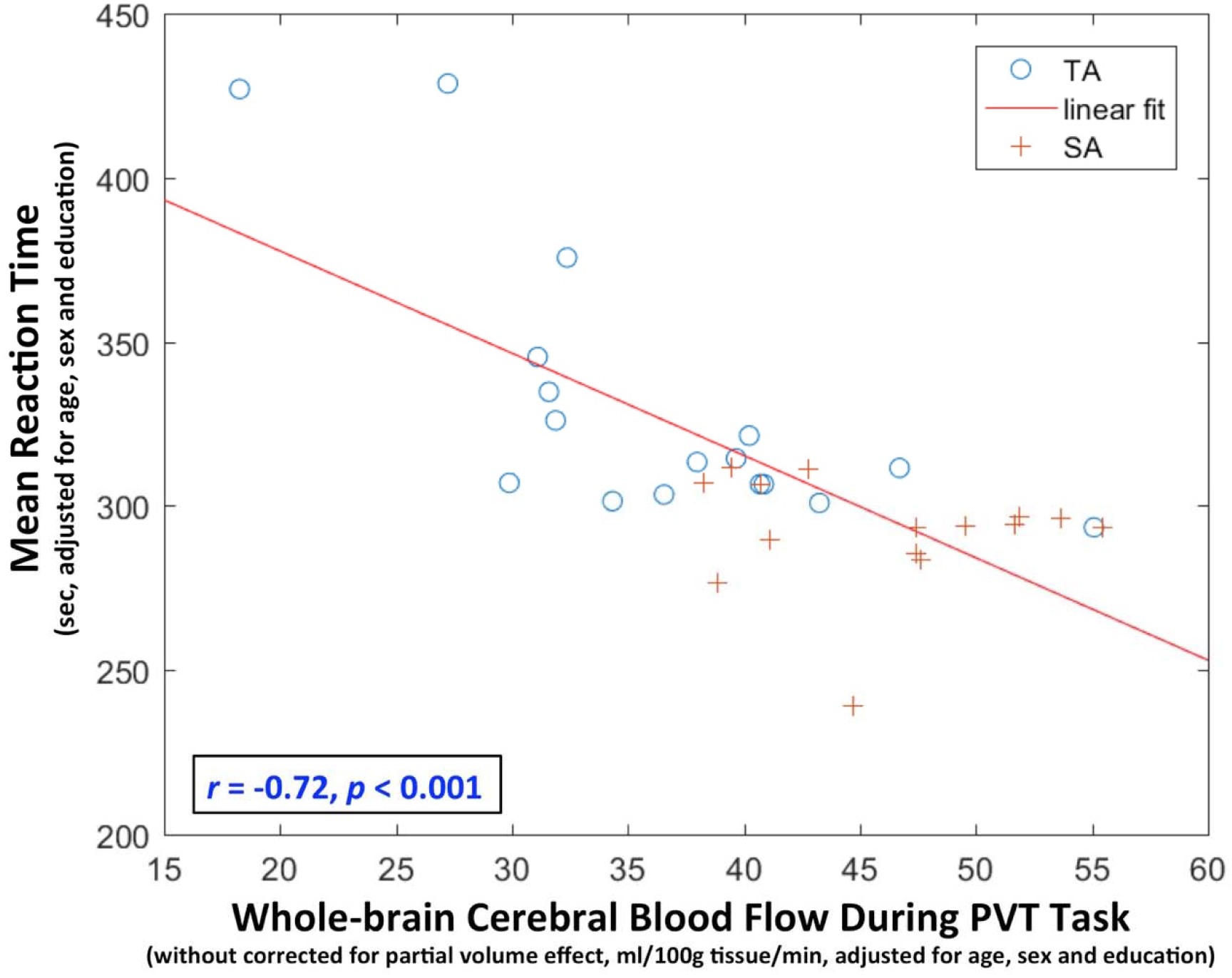
Scatter plot of whole-brain cerebral blood flow during the Psychomotor Vigilance Test (PVT) without correction for partial volume effect and the mean reaction time, adjusted for age, sex and education, of TypicalAgers (TA) and SuperAgers (SA).

### B. Statistical maps of cortical thickness and resting CBF differences between young and older controls

Data-driven regions of interest (ROI) were defined as the regions that exhibit significant age-related differences in cortical thickness and resting CBF between young and older controls. Voxel-wise independent two sample t-tests were performed on the normalized resting CBF maps and cortical thickness maps separately, between young and older subjects using the Randomise package (Winkler, Ridgway, Webster, Smith, & Nichols, 2014). Family-wise error rate (FWE) was applied to correct for multiple comparisons. A significance level of corrected p <= 0.001 is used. As cortical thickness is only defined in cortex, the analysis for the thickness maps was limited to voxels with thickness greater than 0.1mm for all the subjects. The statistical maps for cortical thickness and resting CBF are shown in Figures S2-A and Figure S2-B, respectively. The union regions were defined as ROIs for this study (Figure 1B and Figure S2-C).

**Figure S2.**
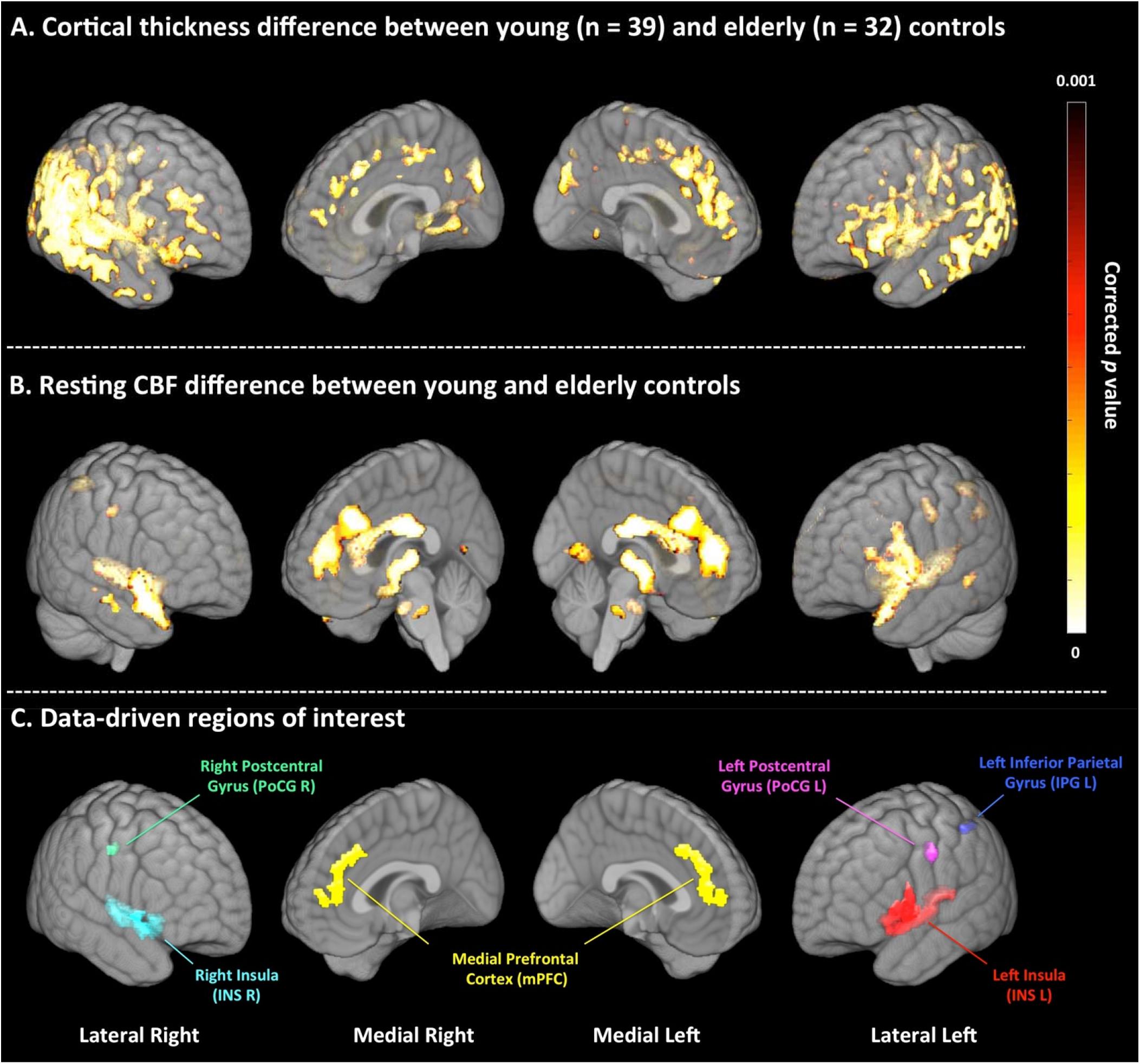
Statistical maps of cortical thickness (A) and resting CBF (B) differences between young and older subjects, tested by voxel-wise independent two-sample t-tests. Family-wise error rate was used to correct for multiple comparisons, with a significance level of corrected p <= 0.001. Regions exhibiting significant differences in both cortical thickness and resting CBF were defined as regions of interest in this study (C). CBF = cerebral blood flow.

### C. Regional and global CBF and cortical thickness measurements of TypcialAgers, SuperAgers and young control

Figure S3 shows the regional and global functional and structural measurements of TypicalAgers, SuperAgers and Young Controls. ANOVA post-hoc analyses were performed to test pairwise group differences. Qualitatively, there is a trend for all the measurements showing that CBF and cortical thickness of young controls is greater than that of SuperAgers, which is greater than that of TypicalAgers. All of the measurements are significantly different between TypicalAgers and young controls. SuperAgers and Young Controls differed on all measurements except whole-brain CBF during the PVT and at rest (indicated by the red circles in Figure S3). Group differences between SuperAgers and TypicalAgers are reported in the Results section of the manuscript.

**Figure S3.**
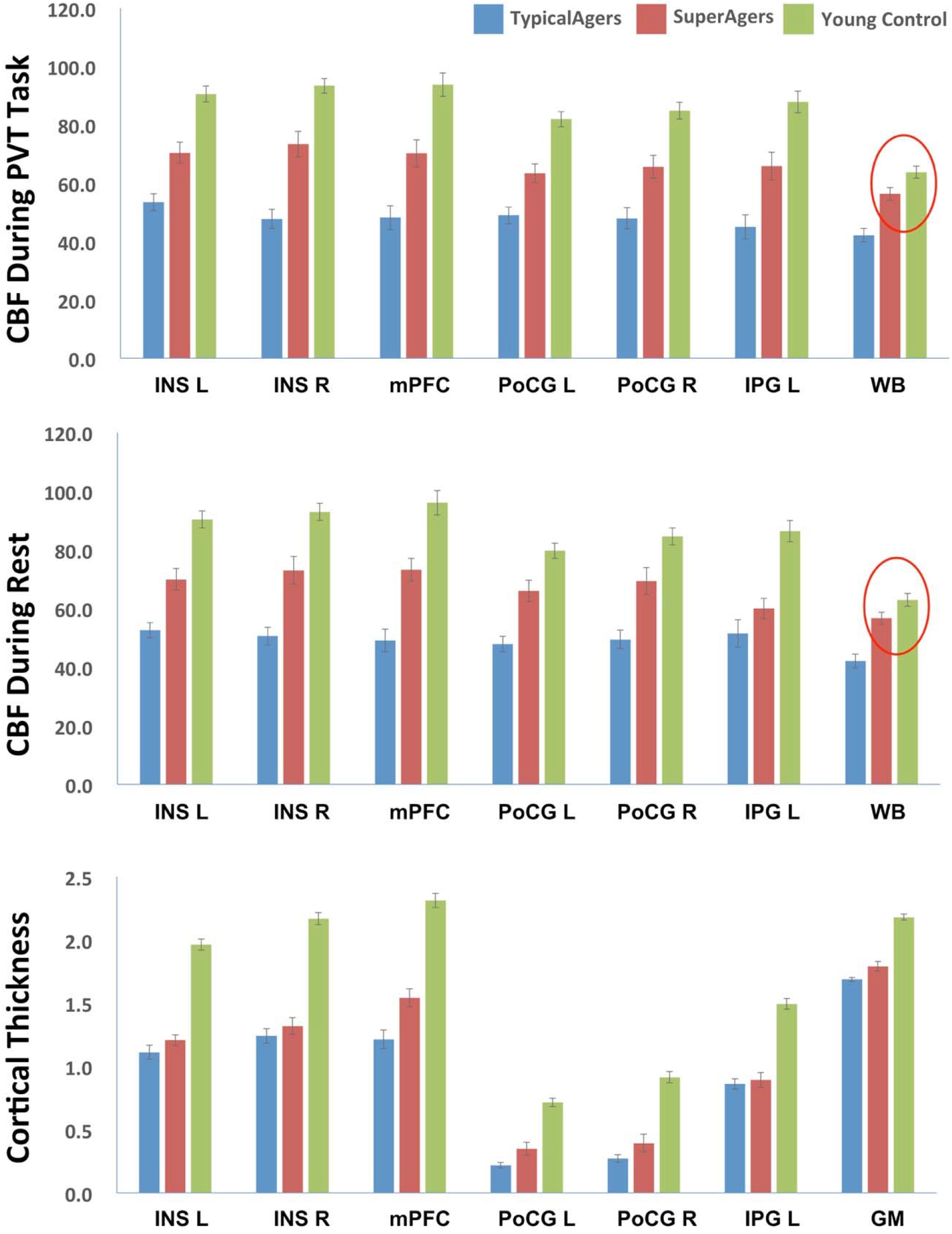
Regional and global cerebral blood flow (CBF) and cortical thickness measurements of TypcialAgers, SuperAgers and young control. Red circles indicate measurements that are not significantly different between Young Control and SuperAgers after Bonferroni correction. INS = insula, mPFC = medial prefrontal cortex, PoCG = postcentral gyrus, IPG = inferior parietal gyrus, WB = whole brain, GM = gray matter, L = left, R = right.

### D. Voxel-wise partial correlation between resting/task CBF and mean reaction time

To investigate whether the significant correlation between global CBF and mean reaction time (RT) is driven by specific brain regions, voxel-wise partial correlation analyses were performed between resting and task CBF and mean RT on the PVT. A general linear model was fit with mean RT as the dependent variable and resting/task CBF as the independent variable, with age, sex and education as covariates. Threshold-free cluster enhancement (TFCE) (Smith & Nichols, 2009) available in the “randomize” package (Winkler et al., 2014) was used to enhance the statistical maps, which were then converted to voxel-wise corrected *p*-values by applying permutation testing with 10,000 iterations followed by family-wise error rate (FWE) correction. A significant level of corrected *p* = 0.05 was used to identify areas with significant prediction value. The results, shown in Figure S4, demonstrate that resting and task CBF of all the brain regions was significantly correlated with mean RT such that greater CBF was associated with lower mean RT, indicative of faster processing speed. Thus, the significant correlation between global CBF and mean RT is likely a whole-brain effect rather than driven by specific brain regions.

**Figure S4.**
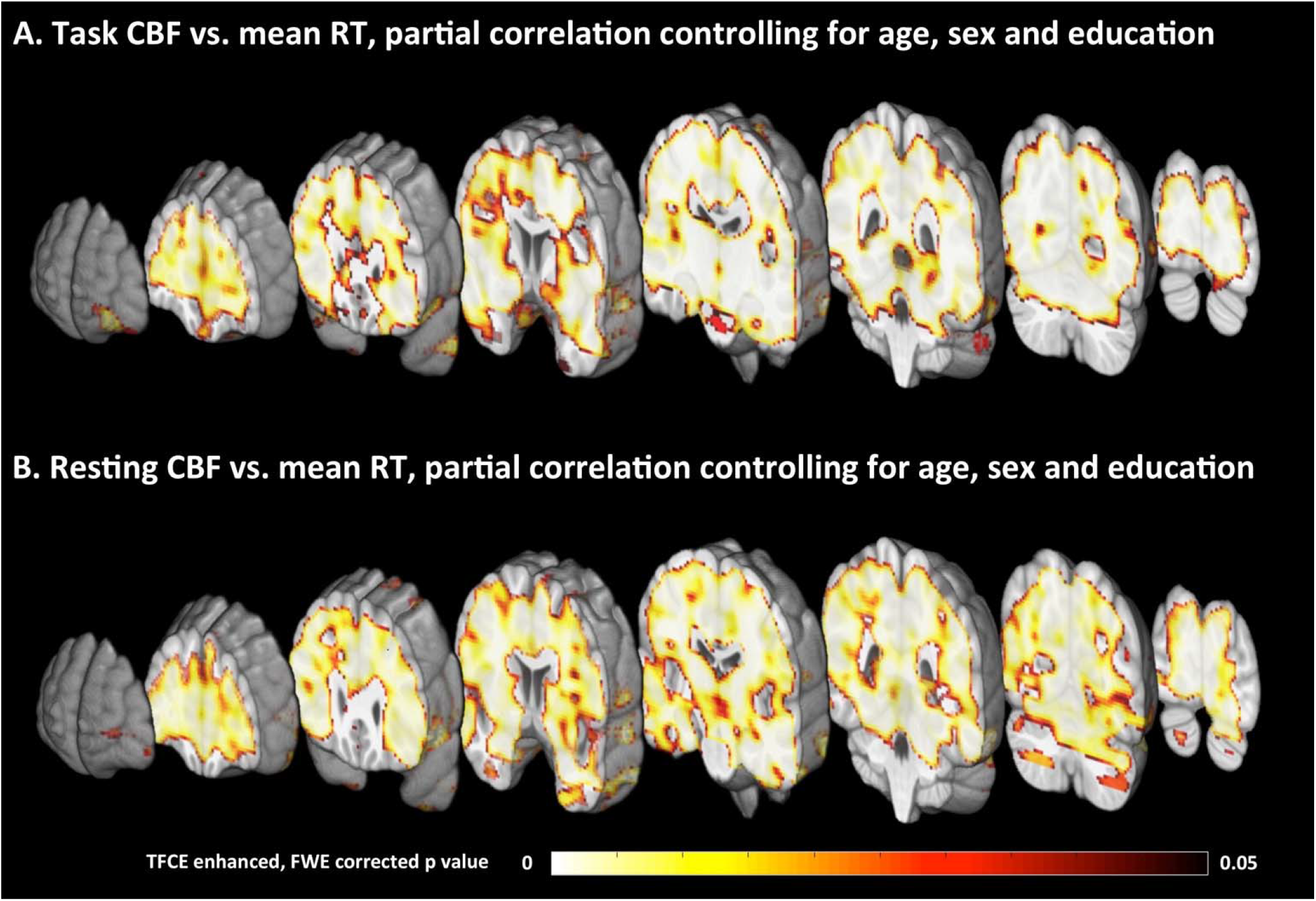
Statistical maps of voxel-wise partial correlation. between CBF during PVT task (A) and resting CBF and mean reaction time (RT), controlling for age, sex and education. TFCE = threshold-free cluster enhancement, FWE = family-wise error rate.

