## Supplementary material for "Preserved global cerebral blood flow accounts for youthful processing speed in older adults": Table S1, Table S2, Figure S1, Figure S2, Figure S3, Figure S4

**Supplementary Materials**

**A. Statistical analysis results using CBF measurements without corrected for partial volume effect.**

All of the statistical analyses were repeated using CBF measurements without correction for a partial volume effect. The results, shown in Table S1, Table S2, and Figure S1 are similar to those with partial volume correction. Similarly, the two-step, hierarchical linear regression using the uncorrected CBF measurements generated similar results with only whole-brain task CBF (*β* = -3.1, *R^2^* change = 0.457, *p* < 0.001) selected in the most predictive model (N = 32, *F* = 7.7, *R^2^* = 0.532, *p* < 0.001).

**Table S1.** Global functional measurements in Young Control, TypicalAgers and SuperAgers without correction for partial volume effect.

|  | **Young Control (YC)** | **Typical Agers (TA)** | **Super Agers (SA)** | **Cohen’s d Between Groups** | | |
| --- | --- | --- | --- | --- | --- | --- |
|  |  |  |  | YC vs. TA | YC vs. SA | SA vs. TA |
| N | 39 | 17 | 15 |  |  |  |
| Whole-Brain Task CBF  (without Partial Volume Correction, ml/100g tissue/min) | 52.5 (10.7) | 35.4 (7.8) | 47.1 (6.9) | **1.72***** | 0.62 | **1.59***** |
| Whole-Brain Resting CBF (without Partial Volume Correction, ml/100g tissue/min) | 51.7 (11.2) | 35.2 (8.0) | 47.3 (6.7) | **1.59***** | 0.51 | **1.62***** |

Note: Mean (standard deviation). *, *** denote *p* < 0.05, *p* < 0.001, respectively (Bonferroni corrected).

**Table S2.** Partial Pearson correlations between global and ROI-based functional measurements without correction for partial volume effect and mean reaction time, with age, sex and education as covariates.

| **Regions of Interest** | **Task CBF** | **Resting CBF** |
| --- | --- | --- |
| INS L | -0.69***^1^ | -0.46*^2^ |
| INS R | -0.62***^1^ | -0.40*^2^ |
| mPFC | -0.51**^1^ | -0.52**^2^ |
| PoCG L | -0.59**^1^ | -0.55**^2^ |
| PoCG R | -0.55**^1^ | -0.52*^2^ |
| IPG L | -0.60**^1^ | -0.51*^2^ |
| Global | -0.72*** | -0.63*** |

Note: *, **, *** denote *p* < 0.05, *p* < 0.01, *p* < 0.001, respectively; red stars show *p* values survived Bonferroni corrections (*p* < 0.05/21). CBF = cerebral blood flow, INS = insula, mPFG = medial prefrontal cortex, PoCG = postcentral gyrus, IPG = inferior parietal gyrus, L = left, R = right.

^1^ not significant after controlling whole-brain CBF during psychomotor vigilance test task

^2^ not significant after controlling whole-brain CBF at rest

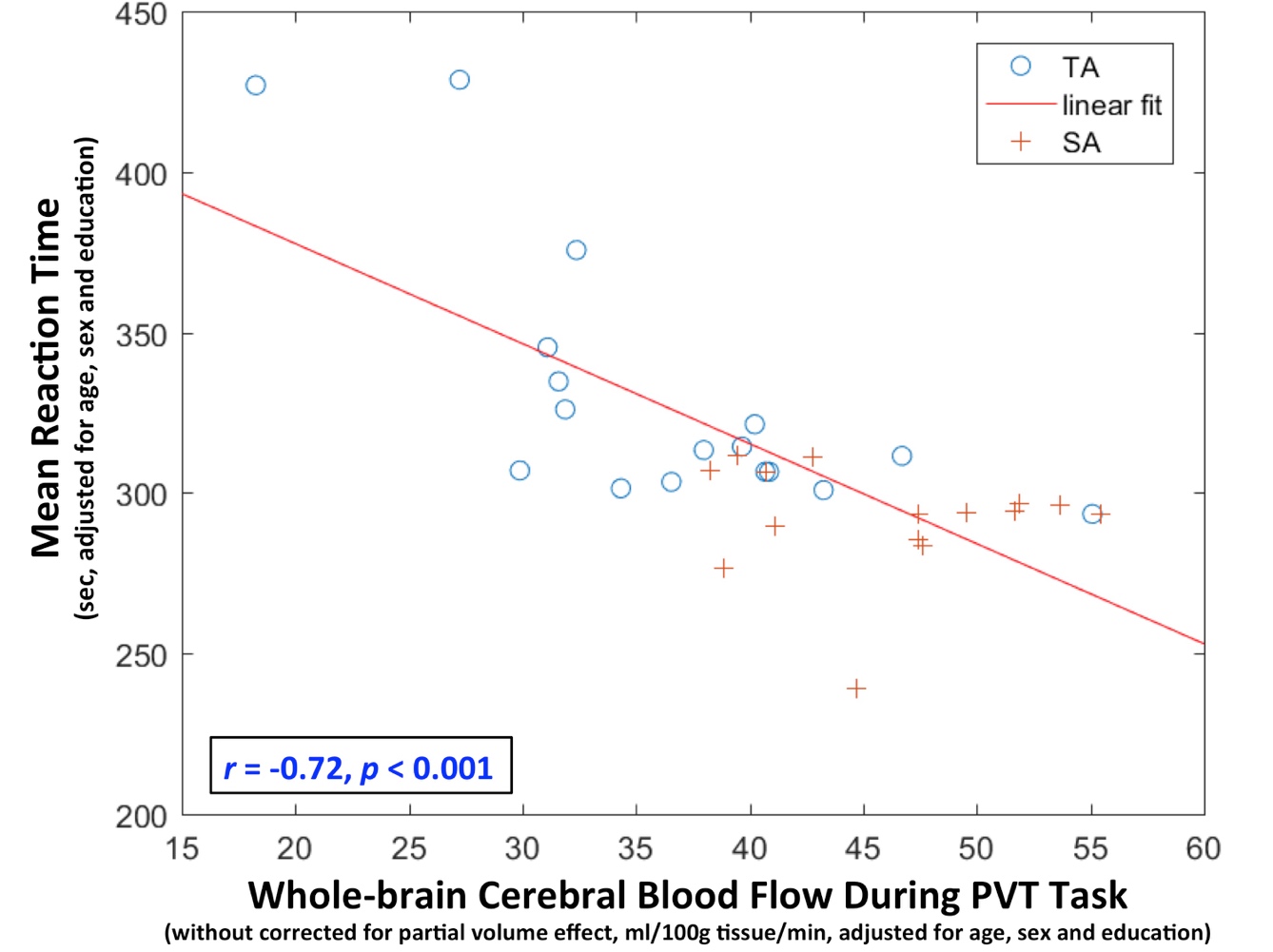

**Figure S1.** Scatter plot of whole-brain cerebral blood flow during the Psychomotor Vigilance Test (PVT) without correction for partial volume effect and the mean reaction time, adjusted for age, sex and education, of TypicalAgers (TA) and SuperAgers (SA).

**B. Statistical maps of cortical thickness and resting CBF differences between young and older controls**

Data-driven regions of interest (ROI) were defined as the regions that exhibit significant age-related differences in cortical thickness and resting CBF between young and older controls. Voxel-wise independent two sample t-tests were performed on the normalized resting CBF maps and cortical thickness maps separately, between young and older subjects using the Randomise package (Winkler, Ridgway, Webster, Smith, & Nichols, 2014). Family-wise error rate (FWE) was applied to correct for multiple comparisons. A significance level of corrected p <= 0.001 is used. As cortical thickness is only defined in cortex, the analysis for the thickness maps was limited to voxels with thickness greater than 0.1mm for all the subjects. The statistical maps for cortical thickness and resting CBF are shown in Figures S2-A and Figure S2-B, respectively. The union regions were defined as ROIs for this study (Figure 1B and Figure S2-C).

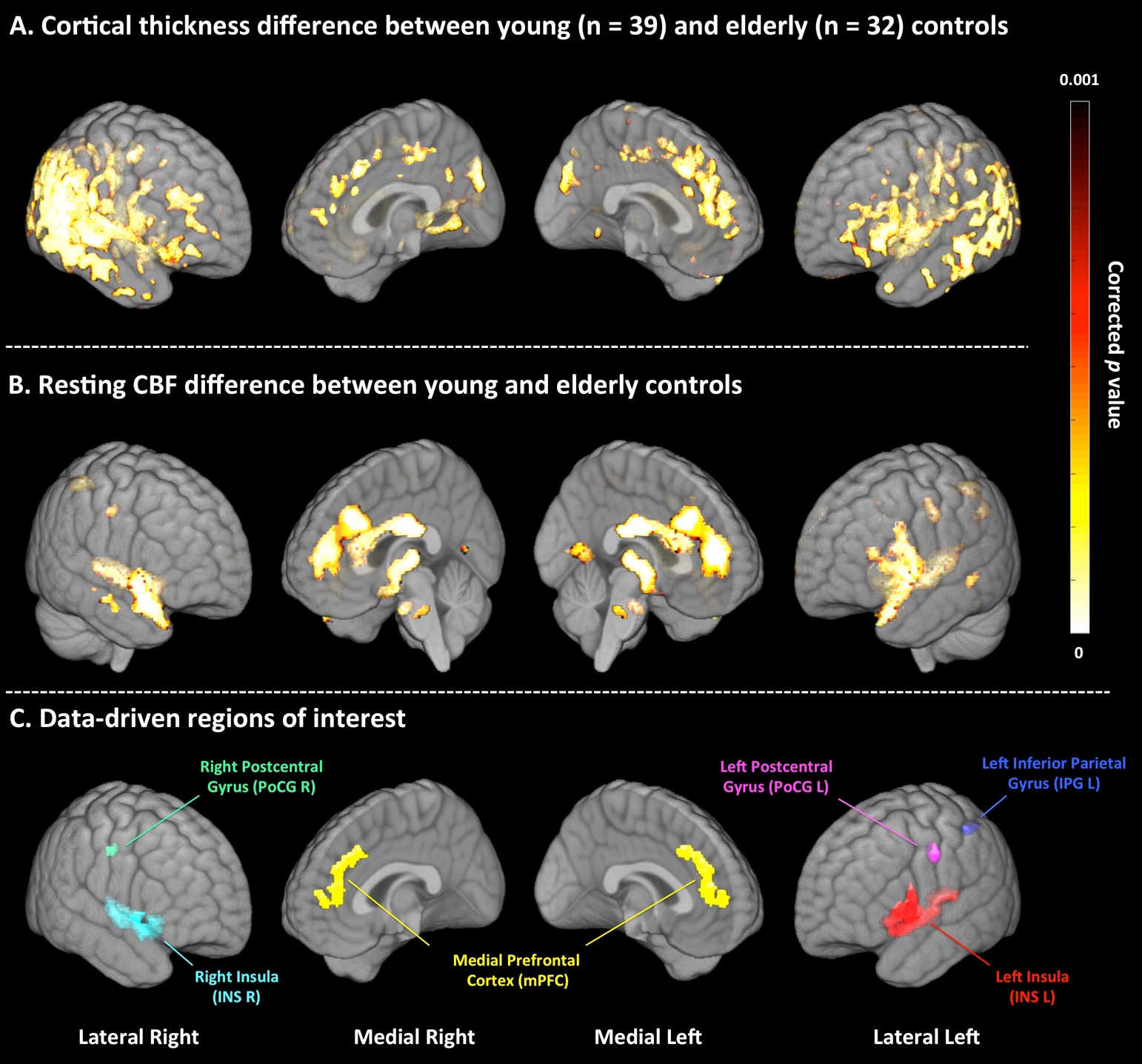

**Figure S2.** Statistical maps of cortical thickness (A) and resting CBF (B) differences between young and older subjects, tested by voxel-wise independent two-sample t-tests. Family-wise error rate was used to correct for multiple comparisons, with a significance level of corrected p <= 0.001. Regions exhibiting significant differences in both cortical thickness and resting CBF were defined as regions of interest in this study (C).

CBF = cerebral blood flow.

**C. Regional and global CBF and cortical thickness measurements of TypcialAgers, SuperAgers and young control.**

Figure S3 shows the regional and global functional and structural measurements of TypicalAgers, SuperAgers and Young Controls. ANOVA post-hoc analyses were performed to test pairwise group differences. Qualitatively, there is a trend for all the measurements showing that CBF and cortical thickness of young controls is greater than that of SuperAgers, which is greater than that of TypicalAgers. All of the measurements are significantly different between TypicalAgers and young controls. SuperAgers and Young Controls differed on all measurements except whole-brain CBF during the PVT and at rest (indicated by the red circles in Figure S3). Group differences between SuperAgers and TypicalAgers are reported in the Results section of the manuscript.

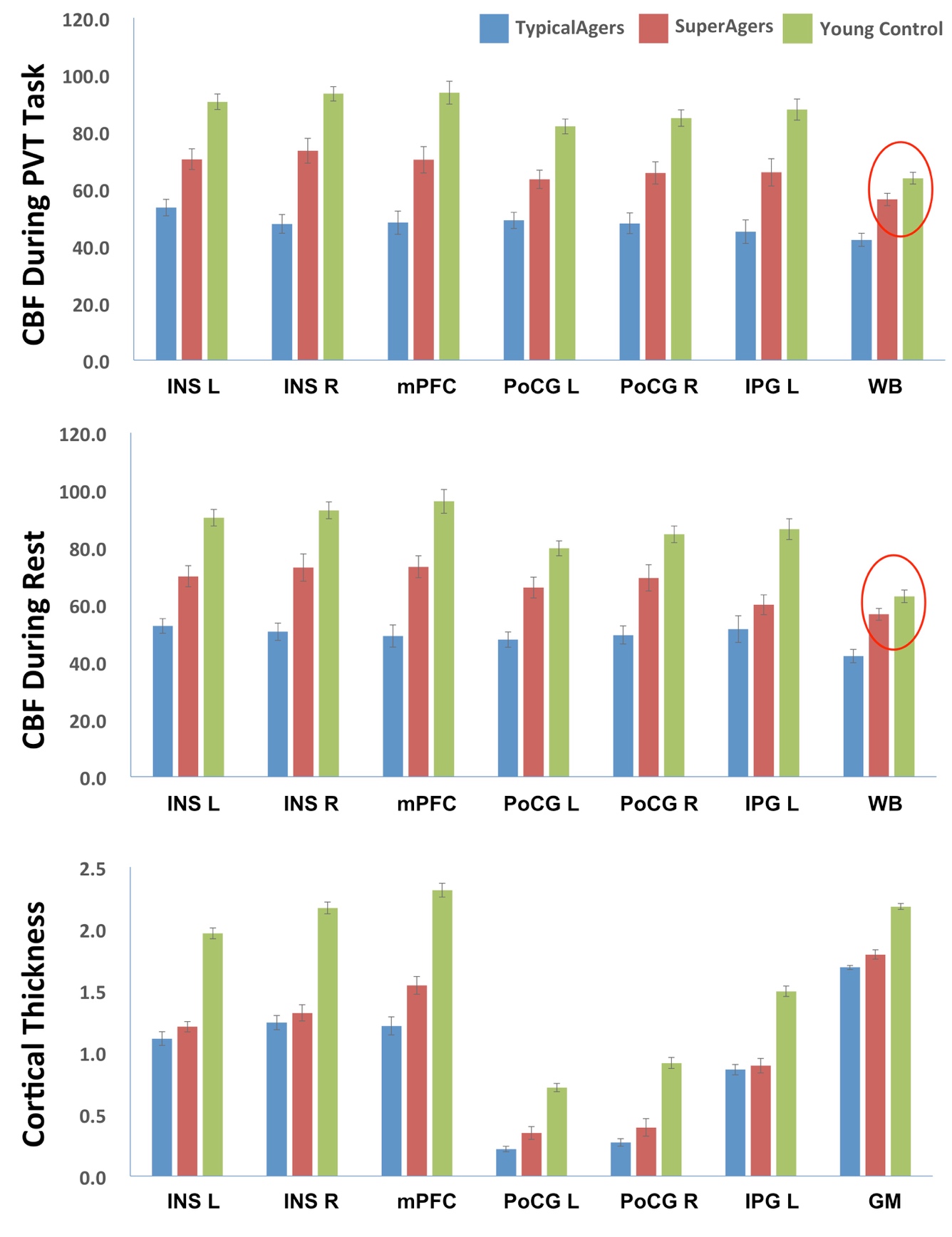

**Figure S3.** Regional and global cerebral blood flow (CBF) and cortical thickness measurements of TypcialAgers, SuperAgers and young control. Red circles indicate measurements that are not significantly different between Young Control and SuperAgers after Bonferroni correction. INS = insula, mPFC = medial prefrontal cortex, PoCG = postcentral gyrus, IPG = inferior parietal gyrus, WB = whole brain, GM = gray matter, L = left, R = right.

**D. Voxel-wise partial correlation between resting/task CBF and mean reaction time**

To investigate whether the significant correlation between global CBF and mean reaction time (RT) is driven by specific brain regions, voxel-wise partial correlation analyses were performed between resting and task CBF and mean RT on the PVT. A general linear model was fit with mean RT as the dependent variable and resting/task CBF as the independent variable, with age, sex and education as covariates. Threshold-free cluster enhancement (TFCE) (Smith & Nichols, 2009) available in the “randomize” package (Winkler et al., 2014) was used to enhance the statistical maps, which were then converted to voxel-wise corrected *p*-values by applying permutation testing with 10,000 iterations followed by family-wise error rate (FWE) correction. A significant level of corrected *p* = 0.05 was used to identify areas with significant prediction value. The results, shown in Figure S4, demonstrate that resting and task CBF of all the brain regions was significantly correlated with mean RT such that greater CBF was associated with lower mean RT, indicative of faster processing speed. Thus, the significant correlation between global CBF and mean RT is likely a whole-brain effect rather than driven by specific brain regions.

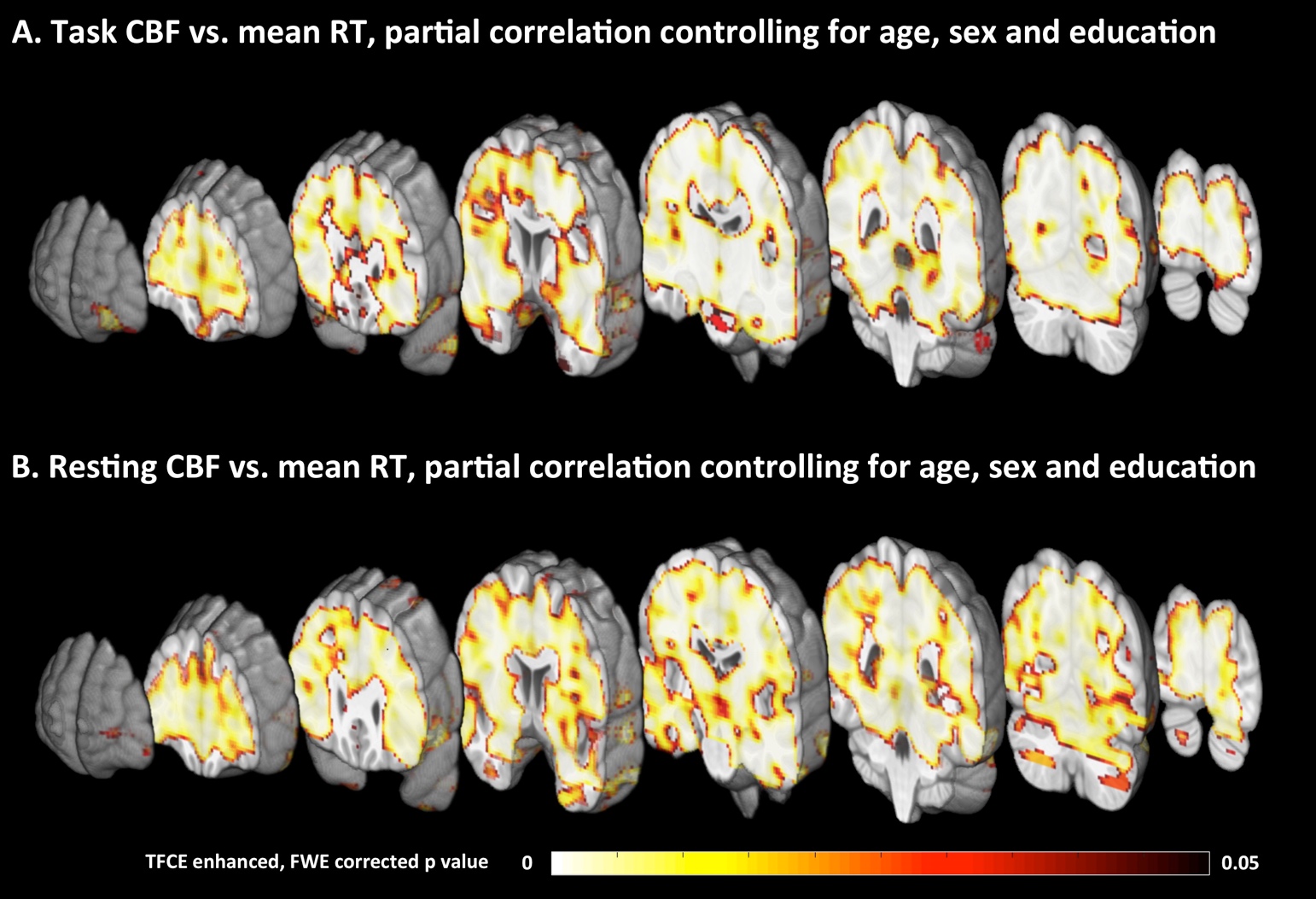

**Figure S4.** Statistical maps of voxel-wise partial correlation. between CBF during PVT task (A) and resting CBF and mean reaction time (RT), controlling for age, sex and education. TFCE = threshold-free cluster enhancement, FWE = family-wise error rate.
